# Ongoing human chromosome end extension revealed by analysis of BioNano and nanopore data

**DOI:** 10.1101/108365

**Authors:** Haojing Shao, Chenxi Zhou, Minh Duc Cao, Lachlan J.M. Coin

## Abstract

The majority of human chromosome ends remain incompletely assembled due to their highly repetitive structure. In this study, we use BioNano data to anchor and extend chromosome ends from two European trios as well as two unrelated Asian genomes. BioNano assembled chromosome ends are structurally divergent from the reference genome, including both missing sequence (10%) and extensions(22%). These extensions are heritable and in some cases divergent between Asian and European samples. Six ninths of the extension sequence in NA12878 can be confirmed and filled by nanopore data. We identify two sequence families in these sequences which have undergone substantial duplication in multiple primate lineages. We show that these sequence families have arisen from progenitor interstitial sequence on the ancestral primate chromosome 7. Comparison of chromosome end sequences from 15 species revealed that chromosome end missing sequence matches the corresponding phylogenetic relationship and revealed a rate of chromosome extension per chromosome of 0.0020 bp per year in average.

## Introduction

The genome sequence of chromosome ends in the reference human genome remains incompletely assembled. In the latest draft of the human genome^1^ 21 out of 48 chromosome ends were incomplete; amongst which five chromosome ends (13p,14p,15p,21p,22p) are completely unknown and the remaining chromosome ends are capped with 10kb-110kb of unknown sequence. There are many interesting observations in the chromosome end regions which remain unexplained, such as the observed increase in genetic divergence between Chimpanzee and Humans towards the chromosome ends^2^.

Chromosome ends contain telomere sequences and subtelomeric regions. Most human chromosome subtelomeric regions are duplications of other chromosome subtelomeric regions arranged in different combinations, referred to as subtelomeric duplications(STD). STD are highly divergent between species or even different populations of the same species^3, 4^ and have experienced rapid adaptive selection^3^. Subtelomere length polymorphism is also found in humans^5^. The majority of subtelomeric duplications have the same orientation towards the chromosome end^3,4^. Based on this it has been suggested that they originated from reciprocal translocation of chromosome tips and unbalanced selection^4^.

Telomere repeat sequences ([TAAGGG]_n_) - which are the capping sequences of chromosome ends - are breakable, acquirable and fusible in the genome. In somatic cells, telomeres are observed to progressively shorten^6,7^ If the telomere sequence is lost, the broken chromosome will become unstable^8–10^, and multiple types of rearrangements can occur, including chromosome fusion^8^, tips translocation^11^, or direct addition of telomere repeats^10^. The manual insertion of telomere sequence in interstitial region results in enhanced chromosome breakages and induces high rates of chromosome rearrangements around the insertion^12^. Interstitial telomeric sequences (ITS) are widespread in the genome^13,14^. In subtelomeric regions, their orientations are almost always towards the terminal end of the chromosome, like the STD^4^.

The quality of assembly for the chromosome ends largely depends on the sequencing technology. To correctly assemble highly duplicated regions like chromosome ends, sequence reads or read pairs spanning the repeat are required^15^. Recently, the NanoChannel Array (Irys System) from BioNano Genomics^16^ was introduced. This technology can generate barcodes on DNA fragments which are hundreds of kilobases long by detecting the distance between specific enzyme recognition sites. Alignment and genome assembly are performed based on numerous distinct site distance fragments. These very long fragments enable construction of individual physical maps, as well as completing the reference in unknown regions^17^. In this manuscript, we report on observed subtelomeric dynamics using this technology for the first time. We conclude that these genome dynamics reflect ongoing chromosome extension and deletion and identify genomic regions which have undergone substantial extension in multiple primate lineages.

## Results

### Assembling chromosome ends using BioNano data

We downloaded data from BioNano Genomics(see Methods). The data contains eight samples, including two family trios (Ashkenazi, CEPH) and two Chinese samples. The raw data has been assembled into contigs (with N50 of 3.3MB) and aligned to the GRCh37 reference. We used the most distal unique aligned sequences at the chromosome ends to anchor individual chromosome ends (see Methods, Figure S1, Table S1-4 and figure 1). We also evaluated this alignment by OMBlast^18^. 92% of terminal contig is still aligned as the same terminal. 7% of terminal contig is unaligned at all with mean size 1.8 Mb. Chromosome X and Y are regarded as one chromosome because their ends are homologous, namely pseudoautosomal region(PAR). We removed 10 terminals from the analysis because of heterochromatin (13p, 14p, 15p, 21p and 22p) and reference gap (1p, 2q, 12p, 17p and XYp). We found that 12%(34/304) and 22% (64/304) of the terminal showed signal of missing and extending sequence, respectively. After removing the two relate samples, missing and extending rate is 11%(29/228) and 21%(47/228). 21 terminals (1q, 2p, 3p, 4p, 5q, 5p, 6q, 8p, 8q, 9p, 10p, 11q, 12q, 13q, 16q, 17q, 18q, 18p, 19p, 21q and XYq) are identical to reference(see Methods). 4 terminals(7q, 14q, 19q and 22q) show individual shortening. Pure shortening could not eliminate the influence of misassembly or misalignment. Thus, we focus on extended terminals. 64 individual chromosome ends in the remaining 11 terminals are extended and could be classified into two categories. Category one (30/64) is extending sequence to its terminal homology distal adjacent sequence (3q, 6p, 7p, 9q(Figure 1a) and 11p). Category two (34/64) is extending sequence to unaligned sequence (4q, 6p, 7p, 9q, 10q, 11p, 15q(Figure 1b), 16p, 20p and 20q). This sequence may further extended to known sequence(2/34).

**Figure 1.**
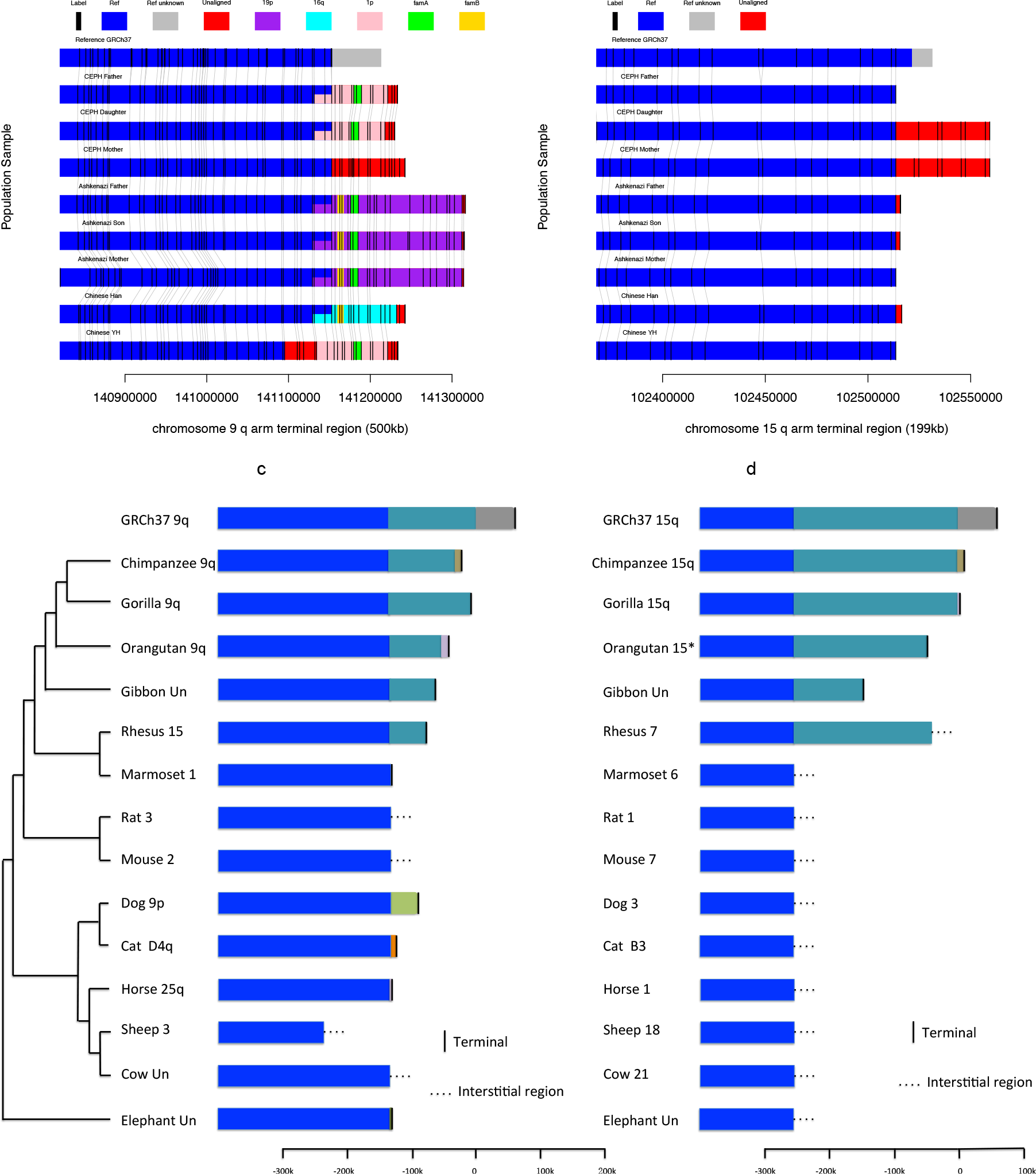
Paralogy map for chromosome tips region. a:) chromosome 9q b:) chromosome 15q for BioNano assembled chromosome ends in eight samples, c:) chromosome 9q and d:) 15q for 14 mammals. The enzyme recognition sites (labels) are marked as black bars and gray connecting lines indicate alignment between samples. Homologous sequence is indicated by color block: blue indicates homology to human reference sequence at the given chromosome; grey indicates unknown reference sequence; purple, cyan and pink indicate homology to 19p, 16q and lp respectively. The remaining unaligned regions are all colored with red. Overlapping colors indicate a shared homology to multiple sources. Yellow and bright green indicate homology with family A and family B respectively. An overlapping color indicates homology two sequences. In c) and d) phylogenetic trees are drawn for the species^19^. The same color indicates the same homology. * means Orangutan chromosome 15 unlocalized scaffold. The solid black line indicates the location of the chromosome end.

The extension sequences supported a re-orientation of chromosome 2q ends in the new GRCh38 reference (Figure S2a). We observed extension sequence at 9q in one family which matched the GRCh38 extension of this chromosome(Figure S2b). We also observed a 260kb extension sequence in one sample at 16p terminal which was originally discovered in 1991^5^(Figure S1.31).

### Discovery of heritable chromosome end polymorphism

We revealed characteristics of chromosome ends by analysis of completed chromosome ends of the eight samples. In the extension terminals, three out of eleven (3q, 7p and 20q) shows almost identical extension sequence, while the remaining eight out of eleven shows population polymorphism. For example, four types of extension sequences (unaligned,1p,19p,16q) are observed in 9q arm amongst 8 samples (Figure 1a). A 45kb extension sequence is observed only in CEPH mother and daughter in 15q(Figure 1b). We observed that the daughter chromosome end sequences (both missing and extending sequence) are almost always (93.8%, 75/80, see Methods) also observed in one of her parents. Although it is a technology limitation to provide a diploid terminal, we manage to identity heterozygous chromosome ends at 4q for two trios(see Methods, Figure S3). The heritable chromosome ends indicate that the chromosome end data are less likely to be a result of an assembly artefact.

### Validating and filling chromosome end extension by nanopore data

The majority of the extension sequence from NA12878 was validated and filled by nanopore data^20^. We downloaded the nanopore data from website(https://github.com/nanopore-wgs-consortium/NA12878). These dataset contains ultra long sequence reads (>100 kb or more), which could use to extend chromosome end and validate the extension sequence(see Methods). Because it is hard to align nanopore data to bionano data directly, we could predict pseudo sequences(see Methods, Table S5) to connect two technology. If the pseudo sequence is well aligned by both data, it is regarded as the true sequence for extension. We could find at least one nanopore read to support six out of nine(3q, 6p, 9q, 15q, 20p and 20q) extension sequence (figure 2). 3q, 6p and 9q extension sequence were 19p, 5q and 1p, respectively. They shared common homologous sequences in the middle(double colour rectangle in figure 2a, 2b, 2c), which were their current terminal ends. 20p and 20q were extended to 1p(with deletion) and 9p with unaligned sequence. 15q were extended to 4 times tandem duplication of its own terminal(about 11kb). Notably, The distal part of 15q is capped with proper telomere. By analysing five nanopore 15q supporting reads, the junction sites of tandem duplication contain different size (0, 900 or 3500 bp) of telomere like tandem repeat sequence(Figure S4, its origin see below). The remaining three extension sequences are uncertain or fail because of weak breakpoint extending(< 15kb) reads (7p and 11p) and unable to predict unknown sequence(4q).

**Figure 2.**
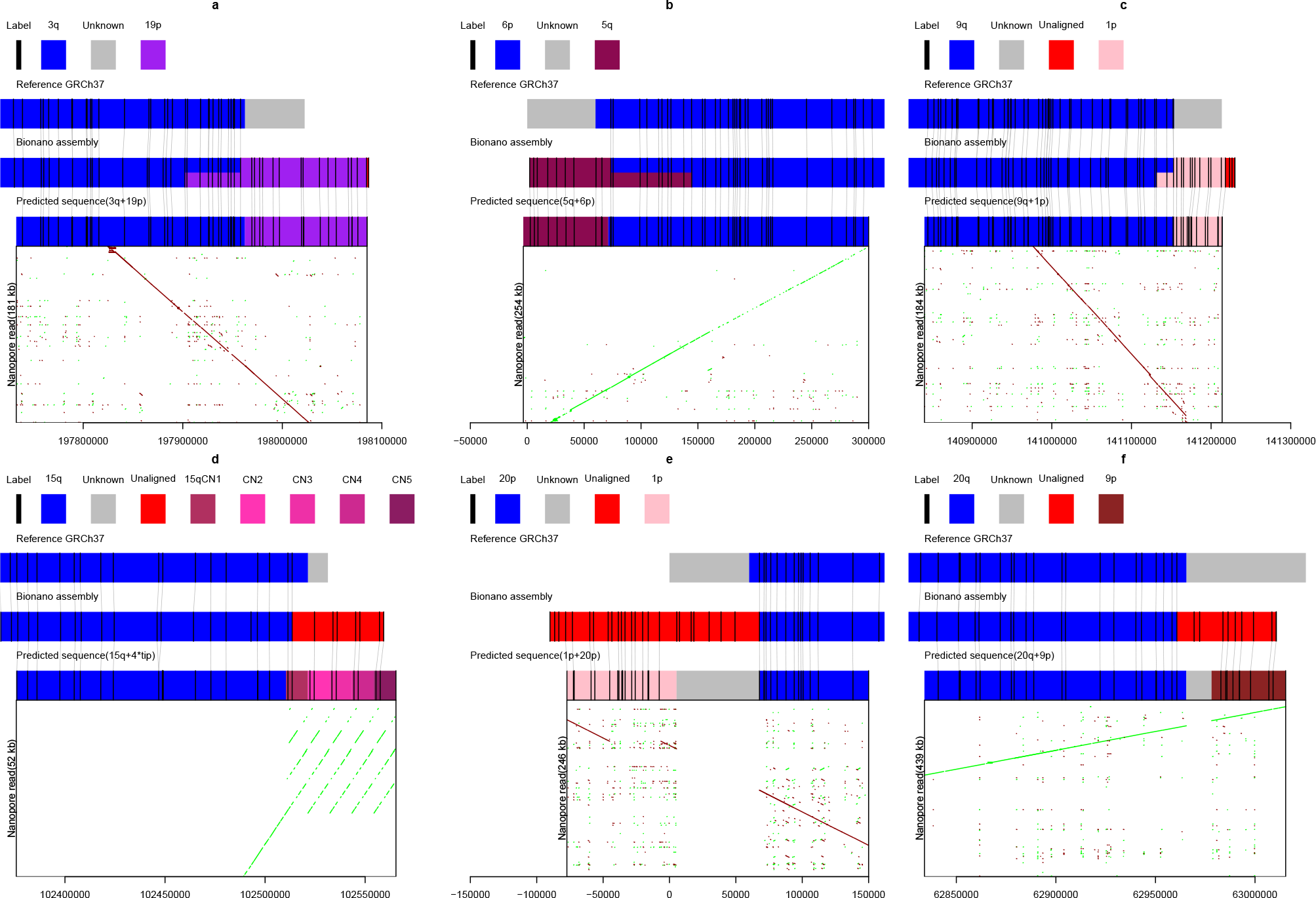
Validation of six terminals by nanopore reads. **a, b, c, d, e** and **f** are for 3q, 6p, 9q, 15q, 20p and 20q, respectively. Reference, bionano assembly in NA12878 and predicted extension sequence(see Methods) are showed as coloured rectangle in the middle. In silico bionano enzyme recognition sites (labels) are showed as vertical black line. The grey lines between labels indicate they are matched. The dotplots of nanopore read to extension sequence are shown at the bottom. Green and red are forward and reverse alignment, respectively.

### Inferring the origin of extension sequences

We recognized the origin of the extension sequences by re-aligning the BioNano labels (enzyme recognition site) patterns to reference. All the aligned extension sequences are aligned to terminal part of 1p, 3q, 5q, 16q or 19p. This indicated some extension sequence was a copy of terminal. These 5 terminal sequences share some homologous sequences with each other. We further checked whether some loci were commonly duplicated during the human extension. We identified (see Methods) two sub-telomeric duplication families (family A and B of length 9kb and 8kb respectively) which were observed in almost all of the 1p, 3q, 5q, 16q, 19p extension sequence regions. We identified all homologous sequences to these two families in the non-redundant nucleotide database which we used to construct a phylogenetic tree for each family(see Methods, Figure 3a,3b, Table S6, S7). These duplications were only found in great ape and were highly duplicated in human (A:16,B:16 copies) and chimpanzee(A:9,B:7 copies). At human subtelomeric regions, both families are clustered within a branch of 100% bootstrap value. The nucleotide divergence between copies are 0.021% to 0.79% and 0.013% to 0.58% for family A and B(see Methods), respectively. Assuming nucleotide substitutions occur at a rate of about 1.5% per 10 million years(3% d vergence between copies)^21^, families A and B duplicated at 0.069 to 2.64 and 0.042 to 1.95 million years ago which were after the divergence of human and chimpanzee. It is consistence with the observation that both families at chimpanzee subtelomeric regions are clustered with themselves. It suggests that the expansion of families A and B occur recently and independently in human and chimpanzee lineage, which also supports the hypothesis of chromosome end extension from duplicating subtelomeric regions. Notable, family A recently duplicated not only at the terminal regions but also at interstitial regions(human Y and chimpanzee 6).

**Figure 3.**
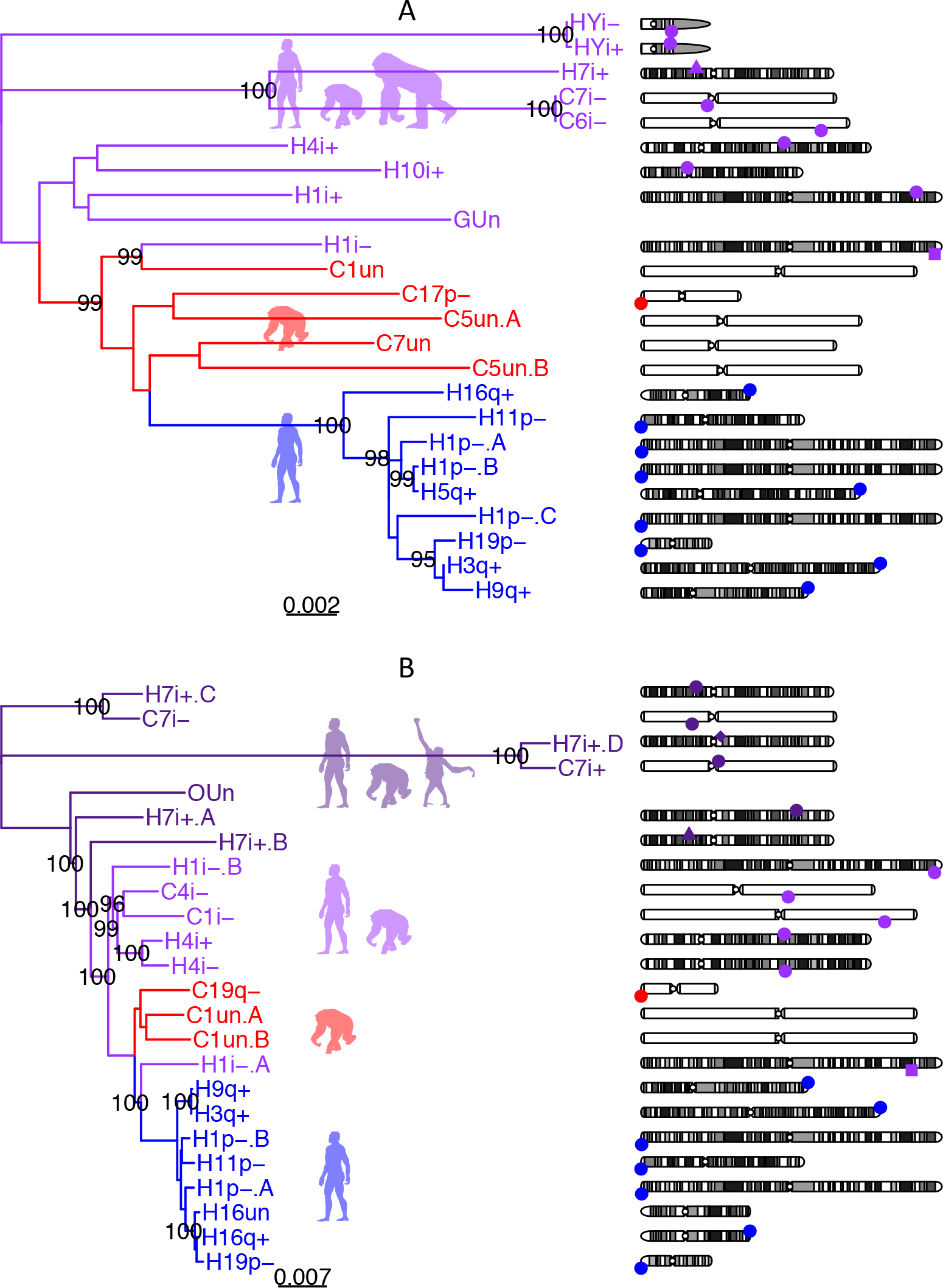
Phylogenetic trees for two sequence familes identified in human extension sequences. A) Family A (9kb) B) Family B (8kb). Phylogenetic trees are generated from their homologous sequences. Dark purple, light purple, red and blue indicate intra-chromosome duplication group, inter-chromosome duplication group, chimpanzee subtelomeric duplication group and human subtelomeric duplication group, respectively. H for Human, C for Chimpanzee, G for Gorilla and O for Orangutan. p for p arm, q for q arm, Un for unplaced contig, un for unlocalised within a chromosome and i for interstitial genome regions. ‘+’ for forward aligned and ‘−’ for reverse aligned. ‘.A’,’.B’,’.C’ and ‘.D’ are appended to distinguish different copies in the same region. Bootstrap value more than 95 are showed. Their genome location are showed as different shapes at each chromosome on the right. The human ancestor copy of intra-chromosome duplication group, inter-chromosome duplication group and subtelomeric group are indicated by the shape of diamond, triangle and square. Otherwise, the shape is circle. Top and bottom indicate different orientation.

A two step duplication history is inferred from the remaining phylogenetic tree. The first step is interstitial intra-chromosome duplication (4 copies) in ancient chromosome 7 in family B(dark purple in figure 3b). The second step is interstitial inter-chromosome duplication from ancient chromosome 7 to other chromosome for both family. Interestingly, these families appeared to both originate from the same region on ancient primate chromosome 7(chr7:45832681-45863525, 7p12.3), separated only by 13kb. In family A, the copy of 7p12.3 achieved highest nucleotide divergence of 4.1%, indicating 7p12.3 was the original copy. In family B, 7p12.3 was not the ancestor copy among other chromosome 7 copy, but the phylogenetics tree(H7i+.B in figure 3b) suggested it was the closest copies to other human inter-chromosome copies (bootstrap value 99.9%), indicating the remaining widespread of family B was originated from this 7p12.3 copy. We aligned to gorilla, orangutan and baboon(see methods) and found that both families were truncated into smaller fragments largely located at ancient chromosome 7. Both families were likely originated from ancient rearrangements.

The ongoing expansion of specific loci at subtelomeric regions observed in human and also reported in chimpanzee and bonobo^22^. They were likely under unknown mechanism. We proposed this process could create new gene, since the duplication of genomic sequences is one of the primary mechanisms for the creation of new genes^21^. For example, the combination of families A,B and other sequences at 11p and 16q create two different genes, namely LINC01001 and FAM157C(Figure S5), respectively. Their expression records could be found at Expression Atlas^23^.

### Identification of the ancient shortening chromosome end telomeric sequence

There are multiple interstitial telomeric sequences (ITS) in the human genome which are orientated in the same direction at subtelomeric regions^4^. For example, 6 telomere sequences are in the first 110kb of 18p (permutation *p* < 0.05, see Methods, Figure 4. We investigated the relationship between all chromosome end ITS and duplications of 1kb or more (Figure 4, Table S8, Methods). All ITS are either fully contained within a duplication, or within 50bp of a duplication (Figure 4d). The vast majority overlap or are next to a duplication on the distal side of the ITS (15 sites) rather than proximal (3 sites, of which each site also has a distal-side duplication overlap, see Table S8). The dominance of distal side ITS duplication suggests that duplication events occurred at the terminal-end of the ITS. We proposed that some current subtelomeric ITS were the ancient chromosome end telomere sequences. Two subtelomeric ITS (chr8:170440-170577, chr19:59097932-59098077) are the most proximal sites of subtelomeric duplication. If no duplication joining to these two sites, these chromosome terminals are unique terminal with telomere sequence like 11q. They are also the start sites where other species could not find any homologous sequence distal to it(8p:7 species and 19q:3 species, see below section).

**Figure 4.**
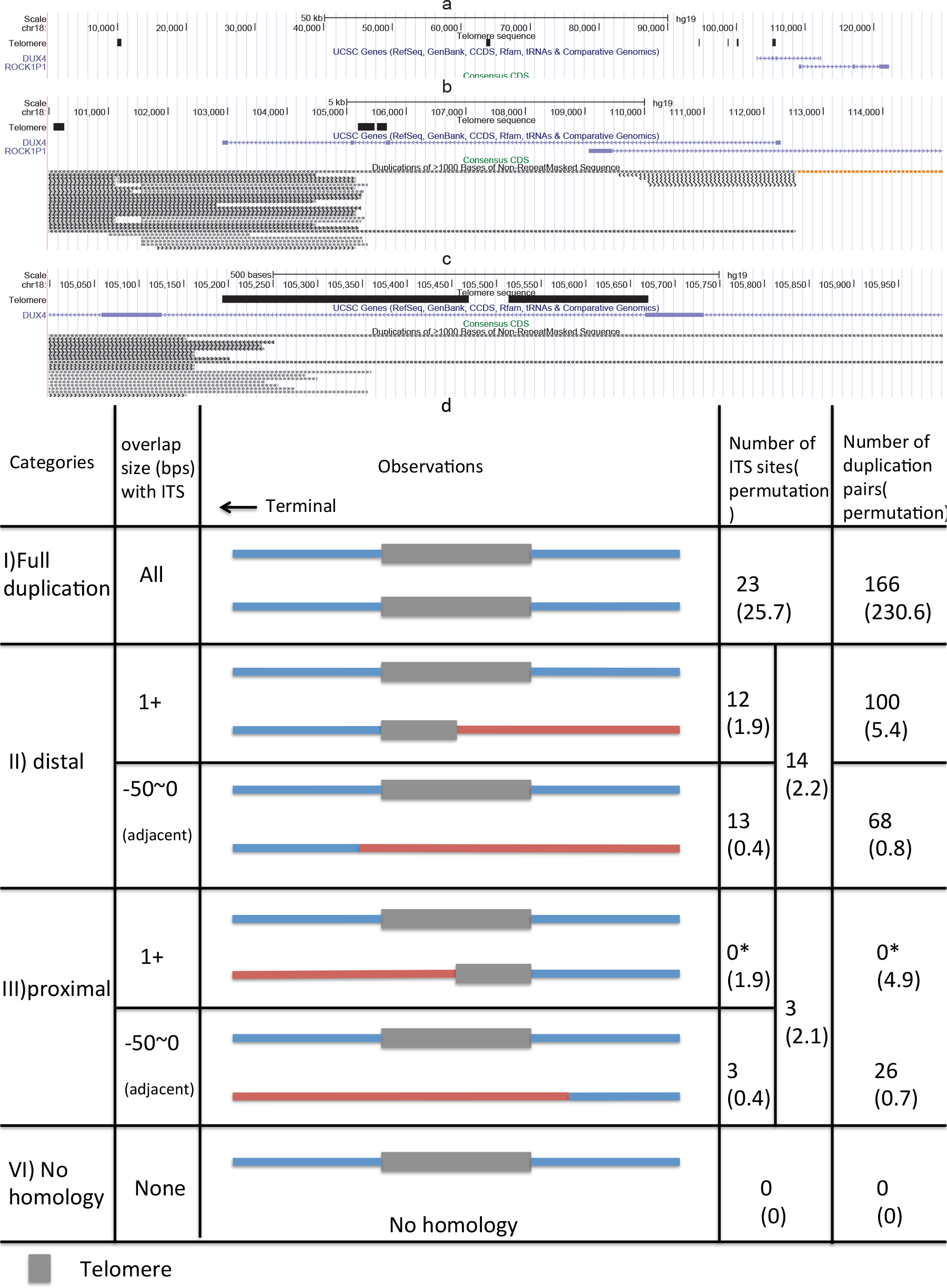
Summary of interstital Telomeric Sequence(ITS). UCSC browser ^25^ displaying chrl8p ITS subtelomeric duplications and genes at three different scales: a) 0-130k; b) 100-114k and c) 105-106k. d) Summary of homology boundary for all subtelomeric ITS. All possible relationships between the sequence containing the ITS (top secontigquence in each cell), and homologous sequence (bottom sequence in each cell) are shown. Blue indicates homologous sequence and red indicates non-homologous sequence. Grey indicates telomere sequence. I) The homologous sequence spans the entire ITS. II) The homologous sequence overlaps the distal breakpoint. III) The homologous sequence is next to (<50bp) ITS distal breakpoint. IV) The homologous sequence is overlapping the proximal breakpoint. V) ITS is next to (<50bp) ITS proximal breakpoint VI) No homologous sequence is observed. * means updating the orientation of 2q and 12p ITS as GRCh38.

We considered three alternate hypothesises for the origin of subtelomeric ITS, but contradictions are found. The first hypothesis is random model, assuming the distribution ITS was mediated by random duplicating or rearranging the subtelomeric sequence. The random permutations find equal random sequences with duplications at both proximal and distal breakpoints (figure 4d), which is significantly different from the observed distribution. The second hypothesis is the reciprocal tips translocation model^4^ for subtelomeric duplication. It does not involve the chromosome terminus, nor create new telomere repeat sequence. The third hypothesis is subtelomeric ITS are insertion sequences of double strand break(DSB) repair. This hypothesis are expanding from Nergadze et al^24^, which is originally proposed for (non subtelomeric) intrachromosomal ITS. The key different between two hypothesises is our model suggests proximal sequence and distal sequence was broken, while ITS insertion model suggests they was joined. We tested two models in human as well as comparison to 5 primates. For all 26 ITS, we checked about all 166 relate duplications(see Methods, Table S9). We found the majority of ITS are the same size with its duplication pairs. Only 2 sites with 5(5/166, 3%) duplication pairs show deletion of ITS. These two deletion(size 1.2 kb, 4.2 kb) contain many other sequences at both proximal and distal site of ITS(size 185, 192 bp). Thus these two ITS are likely deleted with a larger deletion or trimmed by multiple rounds of subtelomeric rearrangement. Comparing to 5 primates, we found zero case that proximal sequence and distal sequence were joined with no ITS(see Methods, Table S10). Distal alignment is still dominant again proximal. There is no proper ITS deleting example to support the hypothesis of subtelomeric interstitial ITS originating from insertion of repairing DSB. And also all alternate hypothesises could not explain the tandem duplication of its own at 15q with telomere sequences(figure 1b). Currently, the most suitable model is that multiple directly duplicated events at ancient chromosome ends have generated multiple present day ITS sequences at subtelomeric regions.

The mean size of subtelomeric ITS (336 bps) is much shorter than capping telomeric sequence. When subtelomeric ITS are used as chromosome end capping telomeres, they create a dysfunctional chromosome end^11^. There are multiple ways to repair the dysfunctional chromosome end^10,11,26^, including chromosome end fusion (Figure 5a), telomere addition (Figure 5b), duplication or translocation of another chromosome end (Figure 5c). Relics can be found for all of these events in the human genome(Figure 5d). Chromosome end fusion is found at ancient chromosome 2A and 2B fusion into chromosome 2^26^, which can be seen from a characteristic inverted interstitial telomere sequence. Telomere addition to telomere is indistinguishable from common telomere shortening and lengthening unless non-telomeric sequence is also involved in the addition. A common observation for ITS or capping telomere is that is has TAR1 (telomere associated repeat 1) element inside, and furthermore the proximal telomere identity is lower than the distal telomere identity(Figure 5d). This suggests that the ancient telomere broke and a new telomere with TAR1 was added. The duplications of other subtelomeric regions to the shortening ITS are the relics of duplication or translocation of other ends to dysfunctional chromosome end. These genome observations are identical to all observations from in-vitro telomere repair models^10,11,26^, suggesting that joining sequence to chromosome ends could occur spontaneously as a result of repairing the dysfunctional chromosome ends both in-vitro, as well as in vivo in our ancestors.

**Figure 5.**
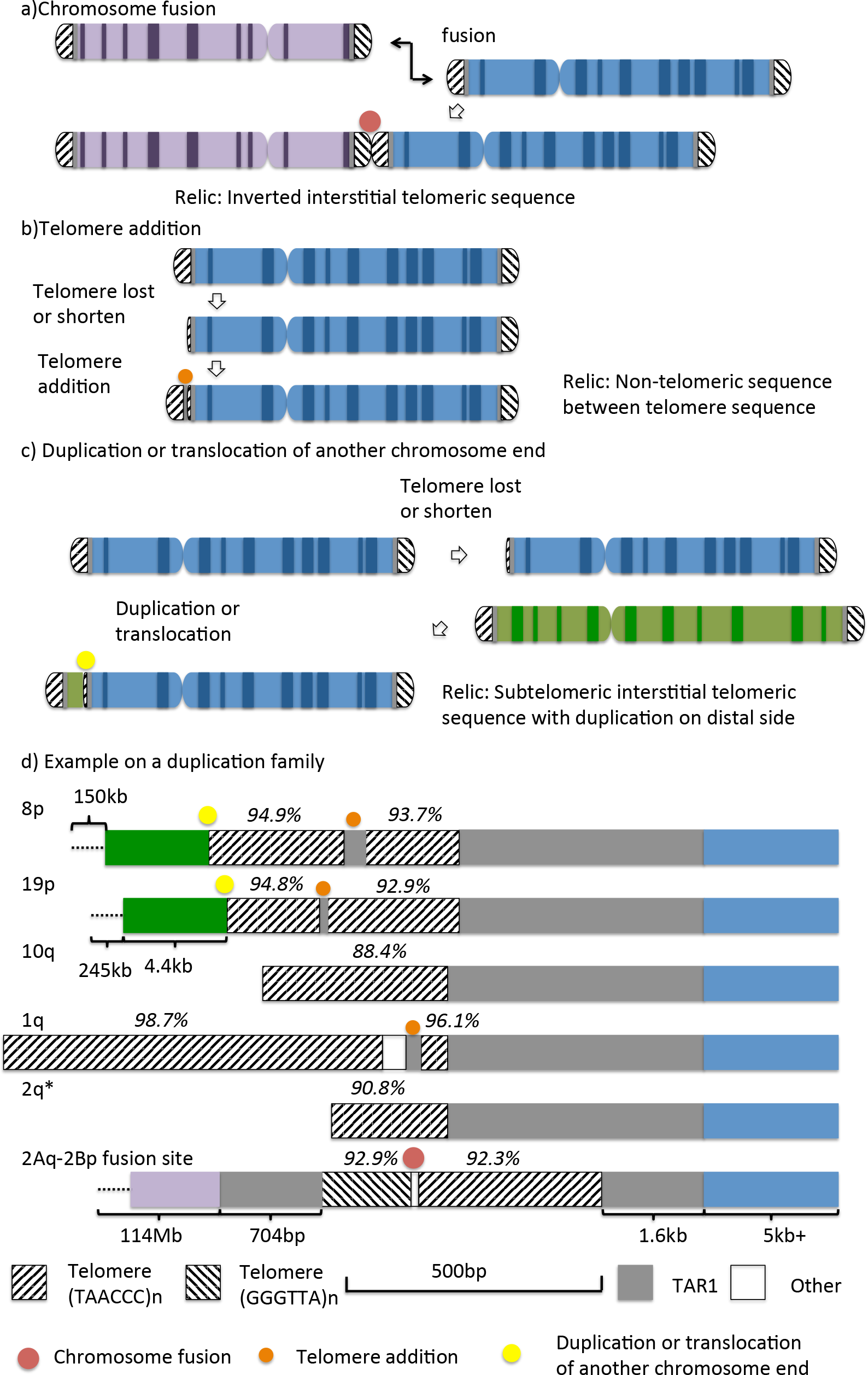
Shortened telomere repair models and example. a. Chromosome fusion, b. Telomere addition, c. Duplication or translocation of other end. d. Example in human genome within one duplication family. The main region is GRCh37:chr8:155249-155739. The size of each block is following the legend except the block with bracket. The telomere repeat identity is showed on the top. * means GRCh38 2q. lOq, lq and 2q are chromosome terminal. The color blocks indicate homology between chromosomes.

### Population genetics at chromosome ends

We used the 1000 Genome Project^27^ data to estimate average genetic diversity at chromosome ends (See methods)[Figure 6]. From these data, we found the diversity decreased (54%) sharply at subtelomeric duplication regions[Figure 6a,6b]. Because these regions may hard to detect SNP and result in lower diversity, we removed the uninformative regions as 1000 Genome Project suggested^27^ and adjusted the estimation(see methods). The diversity is still 15% lower than the adjacent regions. Next, we investigated the divergence of these regions from chimpanzee and found that the divergence was sharply increased at STD(Figure 6c,6d) which was consistence with study^2^. We further found that this increased divergence was mediated by the alignment to paralogous sequence(70%), indicating that chimpanzee missed the homologous sequences(see extension rate section). Combining these observations, the STD regions are special regions in human genome that have lower diversity and high divergence(Figure 6e,6f). The chromosome end extension hypothesis could fully explained these observations. The new extension sequences at STD didn’t have mutations(zero diversity) and took time to accumulate mutation in population(low diversity). And also, these extension sequence didn’t exist in other species, such as chimpanzee. Thus the aligner could only align these sequence to paralogous sequence and result in higher divergence. Otherwise, the divergence should be one or unable to estimate.

**Figure 6.**
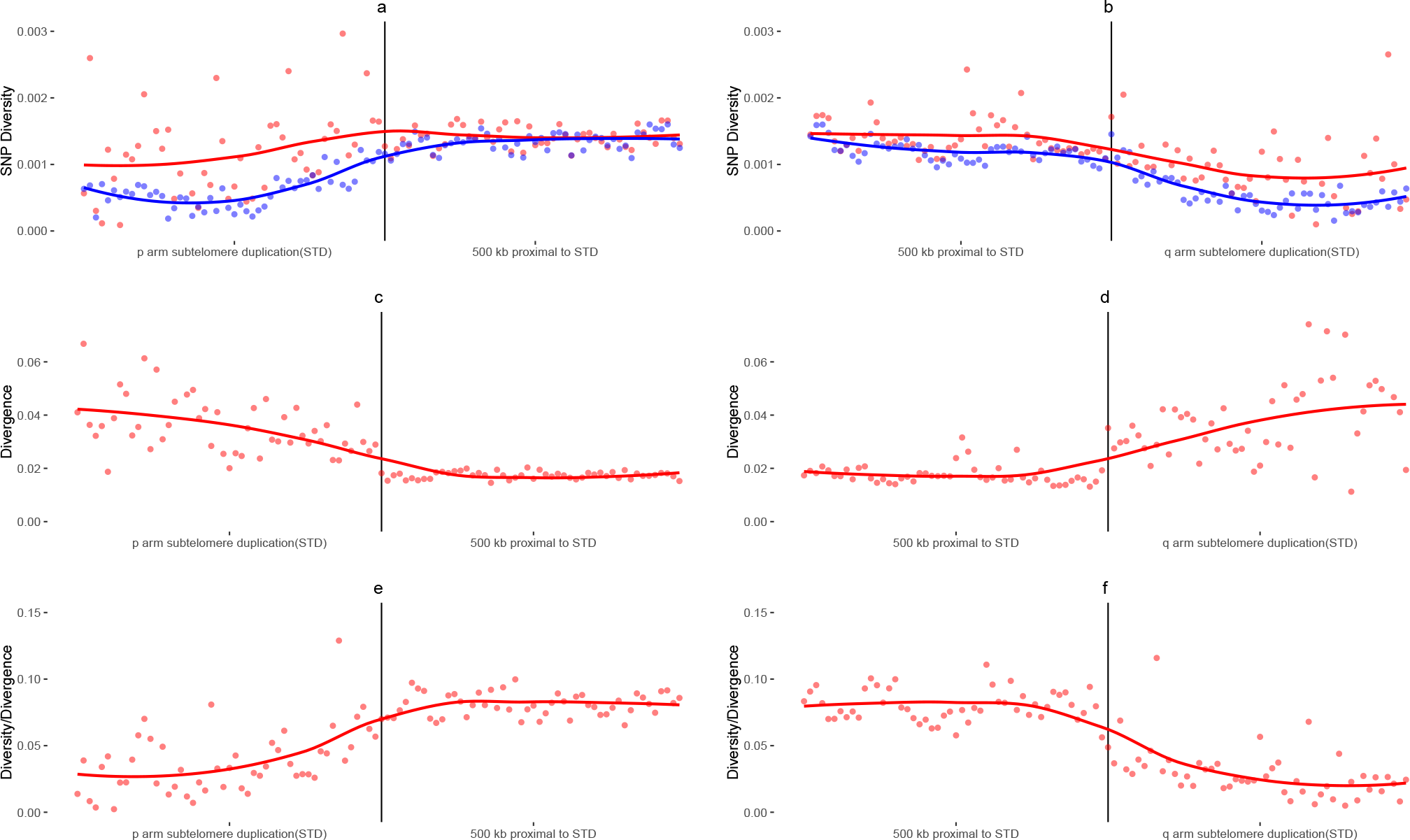
Average population genetic diversity and divergence at chromosome ends. The chromosome end and adjacent sequence is divided into non-overlapping windows. In each window, we calculated diversity(**a** and **b**), divergence(**c** and **d**) and diversity/divergence(**e** and **f**) as dot. Then the line is regression result from the dots. Red and blue are indicated the standard and adjusted estimation of diversity(see methods).

### Human extension rate estimated from species divergence

The comparison of other species chromosome end to human can not only verify the extension hypothesis but also estimate the extension rate. We downloaded pairwise alignments for GRCh37 to 15 well-assembled species from UCSC^25^. 39 well-assembled autosome ends as well as two ancient chromosome 2 fusion ends ^26^ were analyzed (see Methods). By aligning other species to human chromosome ends, we could identify the ancient chromosome end for the most recent common ancestor(MRCA) of human and this species as the most terminal end of the homology and therefore the missing homologous sequence can be defined as human extension sequence since the MCRA (see Methods, Table S11).

Some MRCAs are the same between 15 species, thus their ancient chromosome ends are expected to be identical. We observed that 61% (373/615) of the human-species ancient chromosome ends inferred in this way are identical (Figure 7a and 7b, Figure s6,see Methods, Table S11). In the non primate mammals group, the ancient chromosome ends are highly clustered together, 23 of them are estimated to be identical(labels highlight in Figure 7a,b, see Methods). For example, at 9q and 15q the non-primate mammals are almost all inferred to have the same ends at 134 kb and 255 kb away from the human terminal respectively (Figure 1c,1d). Notably, 50(14%) non-primates mammals autosome ends are still served as current terminals in human(Table S12, see Methods). These chromosome ends contain not only human extension sequence but also other species specific extension sequence, for example, cat D4q, dog 9p and horse 25q(Figure 1c). The extension sequences for human and other species, together with the identical ancient chromosome ends confirm the ongoing extension of chromosome ends.

**Figure 7.**
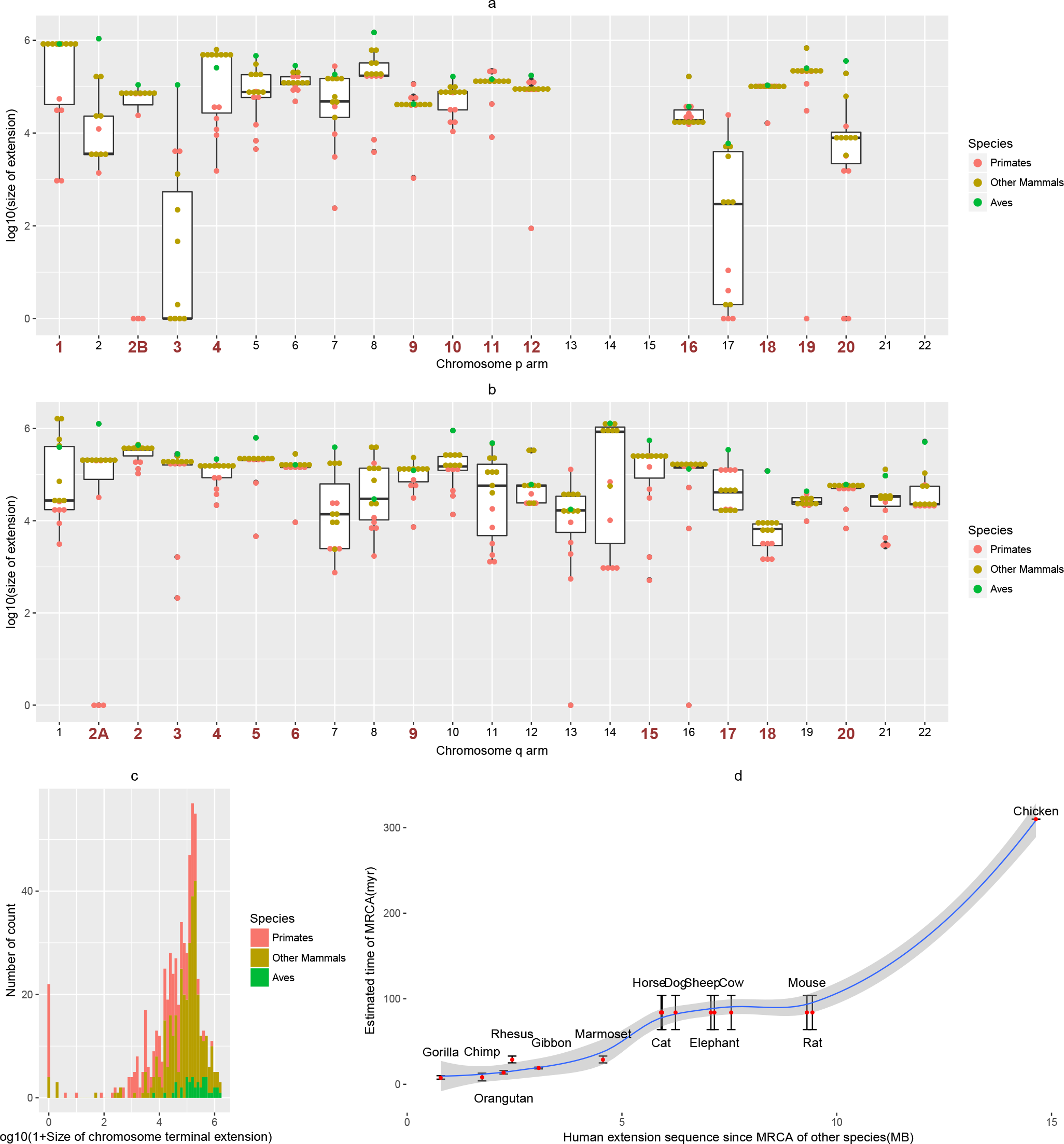
Dotplot, boxplot and size distribution of end extensions. (a) is for p and (b) is for q arm. Each dot is an estimated size of human extension sequence against a species on each arm. Y axis is the normalized size of extension. Bond red numbers indicate these chromosome extension size are clustered together in non-primates group. (c) Length distribution of extension sequence in histogram. (d) Comparison of total extension sequence with MRCA time in 15 species.

The length of human-specific extension sequences represents a near linear relationship with MRCA time^2,19,28–30^(Figure 7c,d), and are consistent with the accepted phylogenetic tree. One exception is that we identified 783 kb of human-specific chromosome extension sequence versus gorilla, whereas we identified 1744 kb versus chimpanzee, which invokes incomplete lineage sorting. However, this may be resolved by the the observation that 30% of the gorilla genome sequence is closer to human or chimpanzee than the latter are to each other^28^.

We also estimated the extension sequence for seven non-primate mammals against each other. Their terminal all contained sequence which couldn’t align by any other species(Figure S7). Their extension size also follow the phylogenetic tree, for example the rat and mouse have less extension sequence compare to each other than to other non-rodent mammals. It suggests the chromosome extension is widespread in mammals.

We estimated the extension rate as dividing the length of extension sequences by the estimated time since the most recent common ancestor (MRCA)(see Methods). This rate estimated the combined effect of both extension and shortening. If shortening is dominant, there will be zero extension sequence like 3p. Considering chromosome end extinction(see discussion), we only estimate the rate in extant chromosome end. In human, we could estimate this rate from the highly identical chromosome ends(count=23, see Methods) which have clear breakpoints among non-primate mammals(Figure 7a,7b). The human extension rate per chromosome terminal since the common ancestor of non-primate mammals is ranging form 0 to 0.0099 bp per year with an average rate of 0.0020 bp per year. The Primates, Rodentia and Eulipotyphla extension rate per chromosome terminal are estimated by comparison to each other. They are estimated to be 0.0021, 0.0036 and 0.0022 bp per terminal per year for 203 Primates, Rodentia and Eulipotyphla, respectively.

## Discussion

In this study we provide evidence that chromosome ends are dynamic and have undergone substantial extension since divergence of primates. We show that the process of chromosome-end extension is ongoing and generates significant polymorphism in the population, which is heritable from one generation to the next.

We identified two duplication families which are participating in ongoing chromosome end expansion. These families appear to both originate from the same region on ancient primate chromosome 7, separated only by 13kb; however they have undergone distinct patterns of duplication in subtelomeric regions of at least two primates (Chimpanzee and Human). These families are still participating in subtelomeric duplication in human populations, as can be seen by alignment of these families to observed BioNano chromosome extension sequence. We showed that duplication of these families in sub-telomeric regions has lead to the formation of new fusion genes. This may provide a mechanism for the extension sequence to be adaptively selected and eventually become fixed in the population. We also showed that formation of extension sequence has resulted in genetic diversity within subtelomeric duplications.

Chromosome end diploid assembly is extremely challenging. For example, NA12878 15q(figure 1b) extension sequence is heterozygous in the sample from the family inherit history. This could also be validated from raw bionano read alignments(11 for reference and 16 for extension). The assembler assumes the non-extended reference type is incomplete sequence and assemble the contig to the longest extended one. The shorter sequence is therefore ignored. Additionally, it is hard to present diploid sequence when the expectation is haploid. The diploid sequence will likely be chopped into small pieces. If the sequence is not complex and long enough, it will output as contig, such as 4q(Figure S3). Otherwise, it is hard to track the diploid sequence. This is the difficulty to determine the zygosity of every chromosome end.

Our analysis indicated that many subtelomeric duplications have been mediated by subtelomeric interstitial telomeric sequence (ITS), and that the duplications occur on the distal side of these ITS. This indicates that the interstitial telomeric sequence are the ancient chromosome ends, and that duplication occurred via a process of fusion to the capping telomere at the chromosome end. Moreover, the observed extensions in the BioNano sequence data appear to be compatible with this hypothesis, although the current resolution of this approach is too large to be conclusive.

The end of human sex chromosome X and Y are identical, namely pseudoautosomal region(PAR). Most of the X short arm was added to ancient chromosome X(X-addition region, XAR)^31^. The PAR could be highly different between species. From the gene scale, the gene inside PAR can be totally deleted or deleted in sex chromosome but locate at autosome, comparing to other species^32^. Sex chromosome addition was often mentioned^31,32^ and an addition-attrition hypothesis was proposed^33^ in 1994. Its mechanism was further hypothesized as terminal interchromosomal rearrangements^32^. The studies of original and evolutionary of X and its terminal(PAR) actually studied the consequence of end extension from chromosome X with an obligatory recombination in large scale. With the new technology, we found sequencing evidences of polymorphism X terminal(PAR) from human population(figure S1.45, S1.46) in smaller scale to support the PAR study^32,33^. The polymorphism autosome terminal suggested the addition was also occurred in autosome. Thus, the chromosome end extension hypothesis is supported by sex chromosome study and further explained in detail.

Considering chromosome X could add a large sequence, we wonder the possibility of larger extension in other chromosome. When the extension become longer(3Mb), the genome likely contains balance copy of the extension sequence. Otherwise, the addition copy of a large sequence in human will likely be negative selected, for example triple chromosome 21. Thus a successful chromosome end large extension may involve chromosome end extinction and creation. As a result, the karyotype may change. It is hard to distinguish with other events(translocation, rearrangement etc.), unless an ancient telomere sequence or other evidences are found. The only example is chromosome 2 fusion^26^, where a pair of proper reverse orientated interstitial telomere sequence is found for ancient 2Aq and 2Bp. This study mainly focuses on small extension. This type of extension could increase the copy of sequence, while larger extension doesn’t contain extra copy and requires a difference method to detect.

Normally, duplication of a subtelomeric region into another subtelomeric regions will never involve interstitial sequence, thus the interstitial origin of sequences at subtelomeric regions indicate an unknown mechanism. Interestingly, two cases of direct interstitial sequence joining to telomere sequence were described in a human cancer fusion study^9^. One was a 447 bps interstitial chromosome 4 sequence joining to 10q telomere at one side and 4p telomere at another side. The other was a 211 bp interstitial chromosome 2 sequence and a 374 bps chromosome 17 sequence joining to 4q telomere and Xp telomere, respectively. These cases in cancer support the ability of interstitial sequence to join to the telomere. A possible model is that the interstitial sequence is deleted and then directly joined to the telomere, similar to the ‘deletion-plus-episome’ model for double minutes in cancer^34,35^.

There are several potential mechanisms we might consider for annealing of chromosome sequence to the capping telomere sequence^4^: non-allelic homologous recombination (NAHR), non-homologous end joining (NHEJ), microhomologous-mediated end joining(MMEJ)^36^ and single strand annealing(SSA)^36^. The presence of overlapping homology at the breakpoints suggests this mechanism is homology mediated. Moreover, the co-location of the breakpoints with interstitial telomere sequence indicates that the process may be partially mediated by micro-homology of the telomeric repeat sequence. Both MMEJ and SSA are also consistent with the observation that all interstitial telomere repeat sequence are in the same orientation.

## Methods

### Bionano data analysis

We downloaded publicly available BioNano data for eight humans (a CEPH trio, an Ashkenazi trio and two Han Chinese) from BioNano Genomics website http://bionanogenomics.com/science/public-datasets/. The initial downloaded data contained raw Bionano data, assembled contigs as well as the unique alignments from contigs to GRCh37 generated using RefAlign from BioNano Genomics IrysSolve. Briefly, this assembler was a custom implementation of the overlap-layout-consensus paradigm with a maximum likelihood model(details see REF^37^). Alignments between consensus genome maps and the hg19 in silico sequence motif map were obtained using a dynamic programming approach where the scoring function was the likelihood of a pair of intervals being similar(details see REF^37,38^). This alignment mapped each Bionano contig to the best matching position on GRCh37, and did not require a full-length alignment of contig to reference, thus allowing us to identify chromosome end polymorphism. We assumed that contigs which are aligned to the most distal sequence with highest matching score represent the individual sample chromosome ends. As the technology is currently unable to resolve different chromosome end haplotypes, the chromosome ends can be viewed as a dominant marker for the longest allele.

The reference chromosome end sequences which could not be aligned by are defined as missing sequences. The reference unknown sequences (”N” region) are removed for missing region size estimation. The unaligned sequence at the distal part of the individual chromosome end is defined as extension sequence. Missing and extension sequences are all more than 33.1 kb(average labels distance+3*standard deviation), otherwise, we defined as the reference type. Chromosomes 13p, 14p, 15p, 21p, 22p were removed because of missing reference sequence. 1p is removed when all the sample’s contigs are discontinuous at the reference unknown region (chr1:471k-521k) and the remaining of 1p could only be aligned as secondly alignment. Sex chromosomes are regarded as one chromosome.

In order to recognize the origin of the extension sequences, we realigned the contigs to GRCh37(373,590 labels) allowing multiple matches by RefAlign. A pre-alignment process in RefAlign was used to merge labels which were close (450 bp) to each other. This process resulted in identification of 346,991 labels for subsequent analysis. The mean distance and standard deviation between adjacent labels was 8.2 kb and 8.3 kb, respectively. The adjacent labels which contain reference gap regions are excluded in estimation. We then used RefAlign to re-align the contigs to the GRCh37 (without chromosome Y) reference enzyme recognition sites (RefAlign parameter: -output-veto-filter_intervals.txt$ -res 2.9 -FP 0.6 -FN 0.06 -sf 0.20 -sd 0.0 -sr 0.01 -extend 1 -outlier 0.0001 -endoutlier 0.001 -PVendoutlier -deltaX 12 -deltaY 12 -hashgen 5 7 2.4 1.5 0.05 5.0 1 1 1 -hash -hashdelta 50 -mres 1e-3 -hashMultiMatch 100 -insertThreads 4 -nosplit 2 -biaswt 0 -T 1e-12 -S -1000 -indel -PVres 2 -rres 0.9 -MaxSE 0.5 -HSDrange 1.0 -outlierBC -AlignRes 2. -outlierExtend 12 24 -Kmax 12 -f -maxmem 128 -BestRef 0 -MultiMatches 5). This re-alignment allowed us to more accurately identify chromosome ends. In particular, for 6p in sample NA12878, the multiple alignments supported a more distal alignment (chr6:73k-364k) than the unique alignment (chr6:245k-1870k). The break of these two contigs was mediated by a connection from 6p to a chromosome 16 interstitial region, which could be an artefact. Similarly, for 16p in NA24385, the multiple alignments supported a more distal alignment (chr16:67k-204k) than the unique alignment (chr16:204k-3728k). The more distal contig could align to both 19p and 16p with no overlap, but the alignment at 19p(chr19: 61k-244k) is longer than 16p making it primarily align to 19p in unique alignment. Notably, the contig with highest matching score for 19p in this sample is another contig(chr19:61k-2341k).

We also realigned individual contigs to other contigs with the same parameter as above. When a label in one sample was aligned to another sample label, we regarded these two labels were connected. For duplication family A and B, we extracted the label IDs from GRCh37 label map. Family A contained 3 to 4 labels. Family B contained 2 labels. For the alignment regions containing family A or B, we only drew these regions when at least 3 or 2 labels were aligned to GRCh37, respectively.

For the CEPH and Ashkenazi trios, we checked the alignments from the child to their parent for every chromosome end(Figure S1). If the child chromosome end was aligned to one of its parent with no more than one label difference at the terminal, we considered that the child chromosome end was inherited from the parent.

We search for potential diploid sequences in the original assembly contigs from realignment. If a sequence from a contig align to two or more locations in the genome, we only consider the location of the highest alignment score as the true location of this sequence. The terminal of 4q in two trios offspring contain two distinguish sequences(see Figure S3).

### Extension validation from nanopore data

Nanopore data from individual NA12878 (rel4) was download form https://github.com/nanopore-wgs-consortium/NA12878. We aligned these data to GRCh37 by minimap2 (https://github.com/lh3/minimap2) with default parameter. We search for the most distal alignments at terminal and identify the terminal supporting reads. Then we search whether the unaligned part of these reads (extension) contain aligned sequence. This analysis enable us to determine the unknown sequence in 15q and 20q as tandem duplication of its terminal and 9p, respectively.

Although the bionano data and nanopore data could not directly alignment to each other, we could predict the sequence and use it to connect two technology. If the extension sequence is aligned in the bionano assembly, it is predicted as connection of the aligned sequence and terminal sequence with estimated gap(”N”). If not, we used the aligned nanopore sequence (15q and 20q) instead if applicable. Otherwise, this extension sequence(4q) is unable to compare and validate. Then we calculate the bionano labels in the predicted extension sequence by software in IrysSolve and aligned them to its assembly(as above, seen as grey link in figure 2). The dotplot was generated from the last^39^(default) alignment of nanopore reads (y-axis) against predicted extension sequence(x-axis). The validation is the correct alignment of both technology to the same sequence.

### Subtelomeric duplication analysis

We downloaded all pairs of regions in the human genome (GRCh37) with high sequence identity from the segmental duplication database^21^. We assign the copy number of a base-pair position to be the number of entries in this file which overlap this position. We take all positions with copy number at least 22 to be high copy number duplication families. We order these families by their base-pair length, defined as the longest contiguous stretch of DNA with copy number greater than 22. Only duplication families which have at least one member within 1MB of a chromosome end, were kept (Table S13). This resulted in an identification of two duplication families with high copy number, which was selected for evolutionary analysis.

We used NCBI blast to align these two sequences (GRCh37,A:chr1:652579-662291,B:chr16:90220392-90228245) to the human genome, chimpanzee genome and nucleotide collection (NT) database, filtering out sequences with less than 95%(human and chimpanzee) or 85%(other) sequence identity to the full query sequence. Next, we extracted the duplications, aligned them by mafft^40^[key option: strategy G-INS-i]. Then we built the maximum likelihood tree by MEGA7^41^ with 1000 bootstrap iterations. Finally, the tree was drawn by online version of TreeDyn^42^. These duplication sequences were also aligned to themselves by lastz^43^(version 1.04.00, parameter default). Then the nucleotide divergence was calculated from the aligned pair(size > 9000 and 7000 for family A and B). Indel were ignored in the estimation. Notably, nucleotide collection(NT) database only contains the sequence of GRCh38, thus we report them in GRCh38 coordinate in this section.

Subtelomeric duplication regions are defined as the longest contiguous duplication sequence starting from the chromosome terminal. From the duplication database^21^, both pairs of duplications regions inside subtelomeric duplication regions are defined as subtelomeric duplications. The longest autosome subtelomeric duplication pairs are chr1:317720-471368 and chr5:180744335-180899425. Homology between the X and Y chromosomes in p arm terminal is maintained by an obligatory recombination in male meiosis^31^. This results in about 2.6 Mb almost 100% identity homologous sequence. Almost all of these sequences are only homologous between X and Y. Since the pattern and mechanism are different between autosome and sex chromosome subtelomeric duplication, sex chromosome is excluded from this analysis.

### Subtelomeric Interstitial telomere sequence analysis

Telomere sequences were annotated from GRCh37 repeatmasker database^44^. We extracted the non-capping telomere sequences inside subtelomeric regions as subtelomeric ITS sequence. We also included the ancient subtelomeric region in the chromosome 2 fusion sites^26^. Then we searched for all the subtelomeric ITS and their adjacent duplications in duplication database^21^. Bases on all the possible overlapping, we divided into 6 types. Type a is duplication spanning the whole ITS. Type b and type d are duplications overlap with the ITS from the distal and proximal site, respectively. Type c and type e are duplications adjacent to (<50 bps) ITS from the distal and proximal sites, respectively. Type f means there is no duplication adjacent to ITS. We counted the number of duplications for each type for each ITS.

We performed 1000 permutations on the ITS. The null distribution is randomly sampling the all the ITS with the same size distribution in the subtelomeric duplication regions. We tested on whether any chromosome terminal distal 121k region contained 5 ITS like 18p. We also calculated the average number of ITS in each category at figure4.

### Chromosome end population genetics analysis

SNP frequencies were extracted from 1000 Genome Project vcf files (v5.20130502). Other mutations are excluded. Chromosome 13, 14, 15, 21, 22 p arm (unknown terminal sequence) and sex chromosomes (different average mutation rate) were excluded from analysis. In the standard diversity estimation, we uniformly divided the subtelomeric duplication regions(definition see above) into 50 windows. Then we uniformly divided sequence 500 kb adjacent to these regions into 50 windows. Considering the difficulty in detecting SNP in duplicated sequences, we downloaded the unmask regions(2.67 Gb in total) from 1000 Genome Project website^27^. We further overlapped this regions with callable divergence region(see below) into merged regions(2.58 Gb). In the adjusted diversity estimation, SNP must inside these merged regions. If a window contained zero merged region, this window was unable to estimate. Otherwise, the diversity in these merged regions will represent the diversity of this window.

For each window, diversity is calculated as the average of the base pair diversity, i.e. 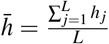, where L is the total (callable) size of the window. The base pair diversity is defined as 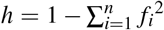, where f is allele frequency and n is the number of observed alleles. Finally, we summarize the average diversity from all the chromosome for each bin as 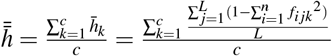, where *c* is total number of available chromosomes.

We downloaded the alignments from human(GRCh37) to chimpanzee(panTro4) at UCSU^25^. In brief, this alignments was unique for each base pair. The divergence regions was defined as the human aligned regions(2.74 Gb). The nucleotide divergence was calculated from all aligned pair. Non SNP mutations were ignored in the estimation.

We performed a local regression on diversity using R(function geom_smooth in ggplot2^45^ package and parameter is method=loess,span=0.7).

### Human extension rate analysis

We downloaded pairwise alignment files from UCSC^25^. These files contain region alignments from species to GRCh37. Initially, 21 genomes (Chicken(galGal3), Chimp(panTro4), Cow(bosTau7), Dog(canFam3), Gibbon(nomLeu1), Gorilla(gorGor3), Horse(equCab2), Marmoset(calJac3), Mouse(mm10), Orangutan(ponAbe2), Rat(rn6), Rhesus(rheMac3), Sheep(oviAri3), Baboon(papHam1), Cat(felCat5), Elephant(loxAfr3), Kangaroo(dipOrd1), Panda(ailMel1), Pig(susScr2), Rabbit(oryCun2) and Zebrafish(danRer10)) were analyzed. For each human autosome as well as ancient chromosome 2A and 2B^26^, we sorted the alignments by human chromosome and location. We searched for the most terminal end alignment. Because short alignments could result from common repeat elements and subtelomeric duplications, we only selected the alignments longer than human longest subtelomeric duplication (154k) to represent ancient chromosome sequence. Because sequence divergence and genome assembly quality will significantly affect the alignment length. Baboon, Kangaroo, Panda, Pig, Rabbit and Zebrafish genomes were hard to represent large ancient chromosome sequence and removed from the analysis.

The human most terminal end of homology are defined as the ancient chromosome ends for the last common ancestor of human and this species. For each ancient chromosome end, we calculated the number of species ends, which has one or more similar ends(within 1kb) with other species end.

The sequence starting from the ancient chromosome end for a last common ancestor to the current human chromosome ends are defined as human chromosome extension sequence since the divergence of human with a given species. Human chromosome end unknown sequences(”N” regions) are removed from size estimation. Total autosome extension sequence (*s*) are the sum of all autosome terminal extension sequences. The human autosome expansion rate since the divergence of human and this species is estimated as 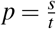, which t is the estimated MRCA time. The average human autosome extension rate since the divergence of human and other primates is estimated as 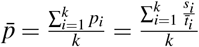, which k is the number of primates and 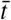 is the mean of estimated MRCA time.

We downloaded the pairwise alignments with each other for eight species (Human(GRCh37), Cow(bosTau7), Dog(canFam3), Horse(equCab2), Mouse(mm10), Rat(rn6), Sheep(oviAri3), Cat(felCat5)) from UCSC^25^ if alignments are available. We also downloaded the unknown sequence annotation files named “gap.txt.gz” from UCSC to infer the terminal unknown sequence(gap). If a species terminal is annotation as “telomere” and there is another gap within 154kb to the terminal, this terminal is regarded as uninformative and removed from analysis like human 13p. Mouse(mm10) chromosome 1 to 19 p arms(3Mb gap with telomere and centromere) and chromosome 4 and 9 q arms(too many gaps) are removed from analysis. For the remaining terminal, we perform a similar analysis like human. Sex chromosome is excluded form this analysis(see discussion).

## Authors’ contributions

H. S. and L.C. conceived the study. H.S. and C.Z. performed the analysis. H.S. wrote the manuscript, which was revised and approved by all authors.

## Acknowledgements

L.C. is an Australian Research Council Future Fellow (FT110100972). The research is supported by funding from the Australian Research Council (DP140103164). H.S. is funded by a University of Queensland scholarship.

